# Development and Validation of a Deep Learning Algorithm and Open-Source Platform for the Automatic Labelling of Motion Capture Markers

**DOI:** 10.1101/2021.02.08.429993

**Authors:** Allison L. Clouthier, Gwyneth B. Ross, Matthew P. Mavor, Isabel Coll, Alistair Boyle, Ryan B. Graham

## Abstract

The purpose of this work was to develop an open-source deep learning-based algorithm for motion capture marker labelling that can be trained on measured or simulated marker trajectories. In the proposed algorithm, a deep neural network including recurrent layers is trained on measured or simulated marker trajectories. Labels are assigned to markers using the Hungarian algorithm and a predefined generic marker set is used to identify and correct mislabeled markers. The algorithm was first trained and tested on measured motion capture data. Then, the algorithm was trained on simulated trajectories and tested on data that included movements not contained in the simulated data set. The ability to improve accuracy using transfer learning to update the neural network weights based on labelled motion capture data was assessed. The effect of occluded and extraneous markers on labelling accuracy was also examined. Labelling accuracy was 99.6% when trained on measured data and 92.8% when trained on simulated trajectories, but could be improved to up to 98.8% through transfer learning. Missing or extraneous markers reduced labelling accuracy, but results were comparable to commercial software. The proposed labelling algorithm can be used to accurately label motion capture data in the presence of missing and extraneous markers and accuracy can be improved as data are collected, labelled, and added to the training set. The algorithm and user interface can reduce the time and manual effort required to label optical motion capture data, particularly for those with limited access to commercial software.

## Introduction

Optical motion capture has been widely used for entertainment, clinical, and research applications to quantify human motion. Passive motion capture systems measure the three-dimensional position of retroreflective markers using a series of infrared cameras. However, these systems are unable to associate the measured coordinates with specific physical markers. For most applications, it is therefore necessary to assign labels to the marker data to define what each marker physically represents (e.g. an anatomical landmark). Given perfect motion capture data, this problem can be formulated as the search for an injective map from a set of three-dimensional coordinates to a set of labels. However, in practice, there may be extraneous markers in the data due to reflection artefacts, known as ghost markers, or undesired markers being in the cameras’ field of view resulting in coordinates that do not correspond to any label. Additionally, physical markers may be occluded at times resulting in labels that are not assigned. The labelling process can be tedious and time-consuming, particularly if the data are noisy and contain occlusions and/or extraneous markers, which occurs frequently in practice.

Previous work has sought to develop automated methods to perform motion capture marker labelling. One approach is to propagate labels forward through time after initializing the labels in the first frame of data (Guerra-Filho, 2005); however, this can fail if the frame rate is not high enough, the marker set is dense, ghost markers are present, or markers become occluded and reappear later. Moreover, any errors that occur are propagated forward through time. Therefore, some have incorporated skeleton models into their procedures to help identify the markers (Maycock et al., 2015; Meyer et al., 2014; Schubert et al., 2015; Yu et al., 2007). These typically require an initialization pose or movement to generate or scale the skeleton model for each participant. Statistical techniques such as graph matching (Xia et al., 2017) and Gaussian mixture models (Alexanderson et al., 2017) have been proposed, but have been sensitive to marker occlusions.

Recent work has sought to apply advances in machine learning to the problem of automated marker labelling. Han et al. (2018) and Rosskamp et al. (2020) rendered the 3D marker coordinates as a depth image and used convolutional neural networks to perform the marker labelling. Ghorbani et al. (2019) framed the problem as the recovery of the correct ranking of a shuffled marker set that could be approached using permutation learning. Jiménez Bascones et al. (2019) used the Adaboost algorithm to select an optimal set of weak classifiers based on geometric relationships between markers to assign marker labels. A limitation of machine learning-based methods is that they require a large amount of existing labelled data to train the model for each marker set. Han et al. (2018) demonstrated the use of synthetic data for model training by generating simulated marker trajectories for their hand marker set from Kinect (Microsoft, Redmond, USA) data for five participants.

Previously proposed marker labelling techniques have reported accuracies ranging from 59.6-99.9%, depending on data quality (Alexanderson et al., 2017; Ghorbani et al., 2019; Han et al., 2018; Jiménez Bascones et al., 2019; Meyer et al., 2014). However, for most users applying these techniques would be a significant challenge as code to implement the methods has not been made publicly available in most cases. Han et al. (2018) shared code providing the architecture of their neural network and some training data for their hand marker set; however, this is only one part of what is required to use the method to label data. There is also commercial software available to semi-automate this process, such as Nexus (Vicon Motion Systems Ltd, Oxford, UK; vicon.com), Qualisys Track Manager (Qualisys, AB, Göteborg, Sweden; qualisys.com), and Cortex (Motion Analysis Corporation, Santa Rosa, USA; motionanalysis.com). These typically require some manual labelling to be performed to initialize the labelling model for each participant. Furthermore, commercial options require the purchase of software licenses and, therefore, may not always be accessible.

Our aim was to develop an open-source algorithm that can automatically label motion capture markers using machine learning in the presence of occluded and extraneous markers. The algorithm can be trained on existing labelled motion capture data or on simulated marker trajectories if there is a lack of labelled training data. Here, we demonstrate both applications and examine the influence of occluded or extraneous markers on labelling accuracy. We also assess the potential to improve the accuracy of the algorithm as data are collected and labelled using transfer learning. We share code and data to generate simulated marker trajectories for any custom marker set, train the algorithm, and perform the labelling as well as a graphical user interface (GUI) that can be used to run the algorithm, visualize the results, and correct any errors.

## Methods

The marker labelling algorithm assigns labels to raw motion capture data based on probabilities predicted using a deep neural network, then verifies and corrects the results based on a pre-defined marker set. We demonstrated the use of the algorithm for two use cases on different data sets. We first trained the algorithm on pre-existing labelled data using Data Set A, a large data set of processed marker trajectories. We then trained the algorithm on simulated marker trajectories and tested the accuracy on Data Set B, which consisted of raw, unprocessed motion capture data for a wide range of movements. With Data Set B, we also showed how performance can be improved as data are labelled and added to the training set using transfer learning. We assessed the accuracy on movements not contained in the training set, examined the impact of occluded and extraneous markers, and compared the results to commercial software.

### Experimental Data Collection

Two motion capture data sets were used. Data Set A included optical motion capture data for 184 participants randomly selected from a larger existing data set (Ross et al., 2018). At the time of testing, participants provided informed written consent for future use of the data and the Health Sciences Research Ethics Board at the University of Ottawa approved the secondary use of the data (H-08-18-1085). Reflective markers were placed at 45 anatomical locations directly on the participant’s skin or clothing. Each participant then performed a series of functional athletic movements including a drop jump and right and left versions of a hop-down, L-hop, lunge, step-down, bird-dog, and T-balance (Ross et al., 2018). Data were recorded at 120Hz using a Raptor-E motion capture system (Motion Analysis Corporation, Santa Rosa, USA). Marker labelling and gap filling were performed using the software Cortex (Motion Analysis Corporation, Santa Rosa, USA). The marker labels were rigorously checked and manually corrected to provide ground truth labels. The gap-filled data with any extraneous markers removed were used for all training and testing as we did not have access to the raw data. These data were randomly divided into a training set of 100 participants (1140 trials), a validation set of 42 participants (474 trials), and a test set of 42 participants (480 trials).

Data Set B consisted of 18 participants performing the same athletic movements listed above as well as a series of functional and transitional movements including three trials of running, one of walking, and three trials each of kneel-to-prone, kneel-to-run, prone-to-run, prone-to-kneel, run-to-prone, and run-to-kneel movements (Mavor et al., 2020). For the walking trial, participants walked in a criss-cross pattern to cover the capture volume. In the running trials, the participants started the trial outside of the capture volume and ran through the volume. For the other trials involving running, the participants ran into or out of the capture volume, depending on the task. All participants provided informed written consent and the protocol was approved by the University of Ottawa Ethics Board (H-06-18-721). In this data set, 63 reflective markers were attached to the participant at anatomical locations and in clusters on rigid, plastic plates. Data were recorded at 240Hz using a Vantage V5 motion capture system (Vicon, Oxford, UK). Unprocessed and unlabelled marker trajectories were exported from Vicon Nexus 2.5 to .c3d files. It was not guaranteed that each ‘marker’ in the .c3d file contained the data for one physical marker and these sometimes contained data for multiple physical markers as they became visible or occluded. Therefore, when the raw data were imported by the algorithm, trajectories were split and added as a new marker trajectory when the distance between the marker before and after an occlusion was greater than the distance to the closest marker in the frame following occlusion.

Data Set B contained extraneous markers beyond the 63 markers that were part of the marker set. Three participants had an additional 23 anatomical markers attached as calibration markers that were not removed for the motion trials. Five participants had the four-marker calibration wand lying on the floor within the capture volume as well as ghost markers resulting from other infrared cameras, and eleven had the calibration wand and infrared cameras and one or two additional markers attached to the participant (C7 and/or scapula). Two versions of Data Set B were created: one with the extraneous markers removed and one where they were retained. The data set was randomly divided in two, with nine participants available to be used for additional training and nine reserved only for testing.

The markers for Data Set B were labelled in Nexus and labels were exported at three stages of the labelling process. The automatic labelling in Nexus uses a participant-specific labelling skeleton. The joint centre locations and marker positions on each segment are estimated based on correctly labelled motion capture data. This is then fit to new motion capture data to automatically assign labels. First, a static calibration trial was manually labelled and used to scale a generic skeleton and calibrate marker locations to create the skeleton for each participant. The automatic labelling pipeline was then run and the resulting labels were exported to produce the first set of labels (Nexus-Stc). Next, a functional range of motion trial was automatically labelled using the statically calibrated skeleton then manually corrected for each participant. These were used to perform the dynamic skeleton calibration in Nexus. This procedure updates the skeleton’s joint and marker locations as well as estimating joint axes of rotation, joint ranges of motion, and expected variability in marker positions. The automatic labelling pipeline was run using the dynamically calibrated participant-specific skeletons for each motion trial and the labels were exported (Nexus-Dyn). Finally, these labels were manually verified and corrected to obtain the ground truth labels for Data Set B. Labelling in Nexus was performed only with extraneous markers retained as they could not be automatically removed from the Vicon input files.

### Simulated Marker Trajectories

We used OpenSim 4.0 (Seth et al., 2018) to generate simulated trajectories for the cluster-based marker set used in Data Set B using the kinematics of 100 participants from Data Set A. First, the 45 markers used in Data Set A were placed on a modified version of the Rajagopal model (Rajagopal et al., 2016). The model was modified to include a neck joint and the range of motion was increased for all joints. Then scaling, inverse kinematics, and a body kinematics analysis were performed in OpenSim for the 100 participants in Data Set A. An OpenSim marker set was created for the 63 markers used in Data Set B and simulated trajectories for these markers were generated based on the body kinematics and local segment coordinates of the markers (Fig. 1). Data were upsampled using cubic spline interpolation to match the 240Hz of Data Set B.

**Figure 1.**
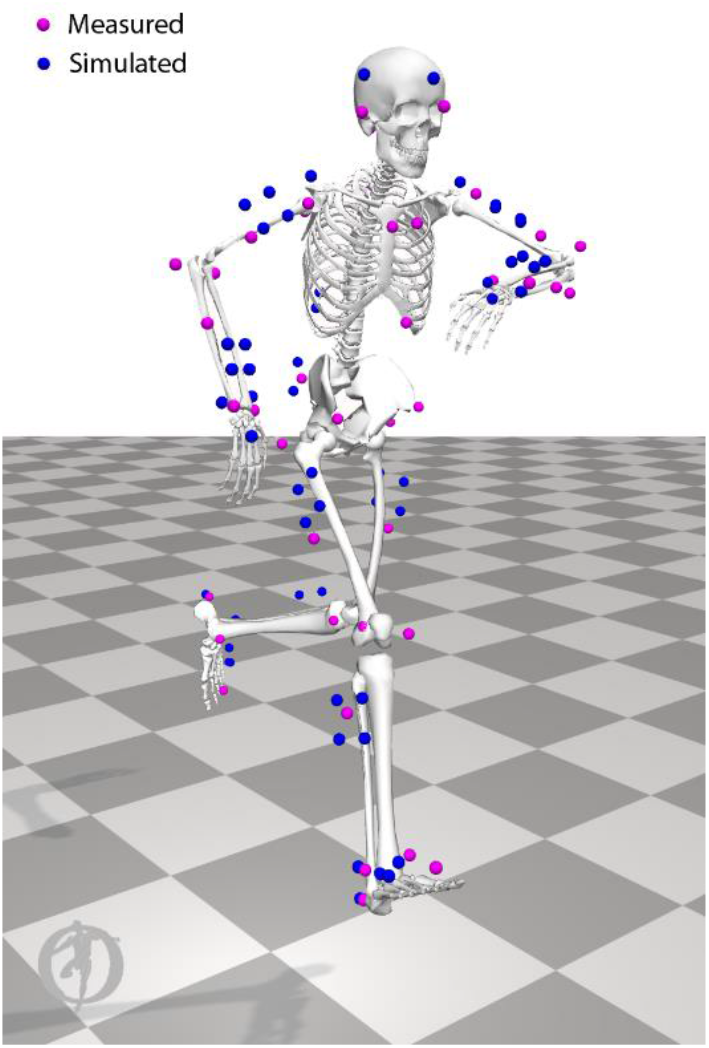
A participant from Data Set A performing an L-Hop. The 45 measured markers are shown in pink and the simulated locations of the 63 markers of the Data Set B marker set are shown in blue.

### Pre-Processing

To prepare the data for input to the neural network, the marker coordinates were first rotated about the vertical axis, so that the participant faced the positive x-direction at the start of the trial. In the training data, this was performed automatically based on the acromion markers. In the test data, the rotation angle to approximately align the data was manually input. Trajectories were low-pass filtered with a zero-phase second-order 6Hz Butterworth filter (Butterworth, 1930) and gaps smaller than 12 frames were linearly interpolated. For each marker, *m_i_*, the marker trajectories were windowed across the visible range of *m_i_*. In the test and validation data, windows of 120 frames were created. If the number of frames was not divisible by 120, the last window would be smaller, unless there were 12 or fewer frames remaining in which case these were added on to the previous window. In the training data, each window was a random length between 12 and 240 frames so that the neural network would have seen windows of different sizes to prepare for the last window of each marker that was usually a different size. If there were more markers than the number of labels, *N_labels_*, then the *N_labels_* markers visible for the greatest portion of the window were retained. If there were greater than *N_labels_* markers with equal visibility, the *N_labels_* markers closest to the marker of interest were retained. Any gaps in the retained marker trajectories were filled with the mean position of the retained markers. The x, y, and z coordinates of the retained markers relative to *m_i_* were calculated and the trajectories were sorted by mean distance to *m_i_*. The Euclidean norm of the velocity and acceleration of the retained markers relative to *m_i_* were calculated. Relative positions, velocities, and accelerations were normalized by the mean values observed across all markers in the training data. A (*window size*) × (5 * (*N_labels_* − 1)) matrix was constructed containing the positions, velocities, and accelerations of the retained markers relative to *m_i_* for input to the neural network (Fig. 2).

**Figure 2.**
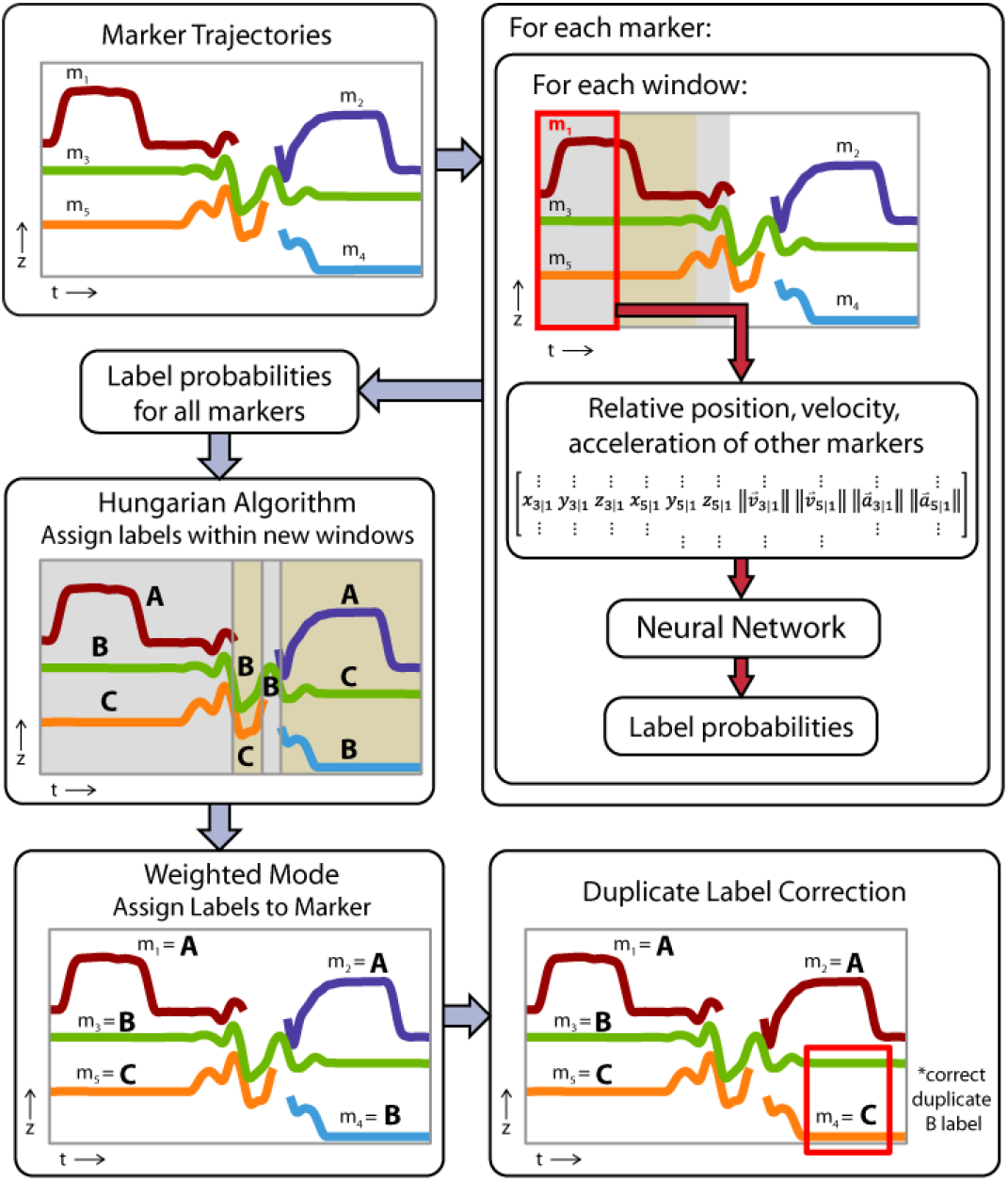
Overview of initial label assignment with an example of three physical markers (labels A, B, and C) that are split into five trajectories due to occlusions.

### Neural Network

A deep neural network was used to calculate the probabilities of the labels for each window of *m_i_*. Five neural network architectures were compared to select the final architecture for the algorithm. These included two architectures using recurrent layers: long short-term memory units (LSTMs) or gated recurrent units (GRUs). These networks included two recurrent layers (LSTM or GRU) with 10% dropout and 128 cells, a fully connected layer with 128 nodes, one-dimensional batch normalization, a rectified linear unit (ReLU), a fully connected layer, and a softmax function. Three established convolutional neural network (CNN) architectures were examined as well, including ResNet-18 (He et al., 2016), ResNet-50 (He et al., 2016), and EfficientNet (Tan and Le, 2019). Each CNN was modified such that the first convolutional layer took one channel as input and the output of the final fully connected layer matched the number of markers in the marker set. All neural networks were implemented in PyTorch (Paszke et al., 2017) and a stochastic gradient descent (SGD) optimizer with momentum was used to train the neural network with a cross-entropy loss criterion. The algorithm was trained for 10 epochs using each of these architectures on the Data Set A training set and tested on the validation set. Training and testing were performed on one NVIDIA Titan RTX GPU on a server with two Intel® Xeon® Gold 6248 CPUs and 384GB of RAM. The accuracy was similar for all networks; however, the training time was significantly less for the recurrent networks (Table 1). Therefore, the architecture based on LSTMs was selected to use going forward.

**Table 1:** Comparison of neural network architectures. Accuracy of the labelling on the Data Set A validation set using various neural network architectures and time to train the neural networks on the Data Set A training set.

|  | Accuracy (%) | Training time (h) |
| --- | --- | --- |
| LSTM | **99.58** | **5.1** |
| GRU | **99.58** | **5.1** |
| ResNet-18 | 99.55 | 31.2 |
| ResNet-50 | 99.57 | 85.2 |
| EfficientNet | 99.55 | 18.4 |

Hyperparameter tuning was performed to select the number of LSTM layers, number of LSTM cells, LSTM dropout percentage, number of nodes in the fully connected layers, and the momentum and learning rate of the optimizer. Optimal parameters were determined through Bayesian hyperparameter optimization using the Ax Platform (Bakshy et al., 2018). The micro-averaged F1 score, which is the harmonic mean of precision and recall and is equivalent to the classification accuracy, was used as the criterion for the optimization. The 100-participant training set and 42-participant validation set of Data Set A were used for hyperparameter optimization. Through this analysis, we found the best results with 3 LSTM layers, 256 LSTM cells, 0.17 dropout, 128 fully connected nodes, a learning rate of 0.078, and momentum of 0.65. The final architecture used is shown in Fig. 3.

**Figure 3.**
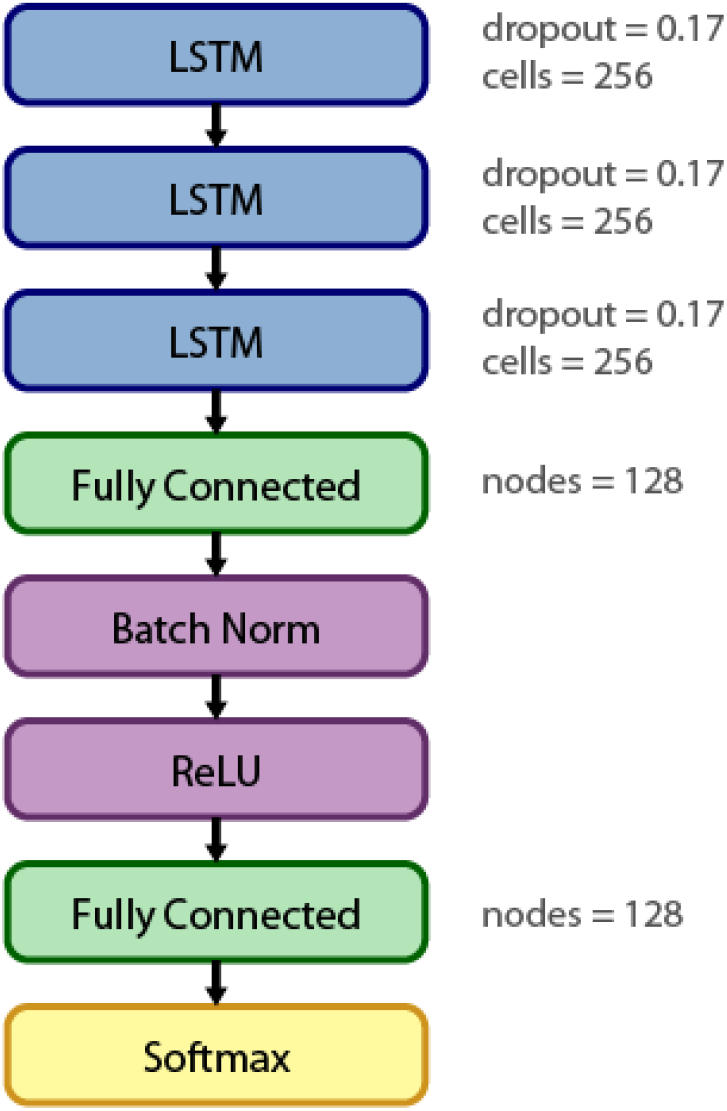
Final architecture of the deep neural network used for the labelling algorithm.

### Label Assignment

The output of the neural network was a 1 × *N_labels_* vector for each window of *m_i_* containing the probabilities of each label being correct. New windows were generated by segmenting the trial marker data at frames where any marker appears or disappears. This ensures that each marker label exists only once in each window, which is necessary because a given marker label may be split across multiple trajectories and this is, therefore, not guaranteed with arbitrarily segmented windows. Assigning labels can be formulated as an unbalanced assignment problem on a weighted barpartite graph. Therefore, for each window, the optimal marker labels were assigned using the Hungarian algorithm (Kuhn, 1955), which has a time complexity of *O*(*n*^3^), to minimize the cost function

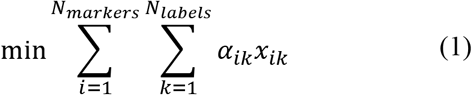

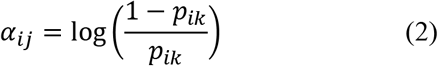

where *x_ik_* = 1 if marker *i* is assigned label *k* and 0 otherwise and *p_ik_* is the probability of marker *i* being label *k* in this window, as determined by the neural network.

At this point, it is possible that windows from the same marker have been assigned different labels. Therefore, the weighted mode was used to assign one label to the entire trajectory. The predicted labels in each frame were weighted by the probability for the prediction. A check was then performed to identify cases where two simultaneously visible markers were assigned the same label and this duplication was corrected by removing the labels from all but the marker with the highest probability.

### Marker Set Verification

The OpenSim (Seth et al., 2018) marker set, which defines the local coordinates of each marker with respect to its associated body segment, was used to identify incorrectly labelled markers based on the assumption that each body segment is approximately rigid (Fig. 4). First, the distances between a marker, *i*, and the other markers attached to the same body segment, *j* ∈ *B_i_*, were calculated and compared to the distances observed in the training data set, *δ_ij_*. If the distances to all other markers on the same segment were outside of three standard deviations of the distances in the training set for that marker, the assigned label was removed. An attempt was then made to assign a label to the unlabelled markers. For each unlabelled marker, labels that had not been assigned during the marker’s visible frames were identified and the mean probabilities were compared. Moving from highest to lowest probability labels, the distances to other markers in the body segment that would result from each available label were calculated and the label was assigned if the distance fell within three standard deviations of that observed in the training set. If the distances were outside of the range for all available labels, this marker remained unlabelled.

**Figure 4.**
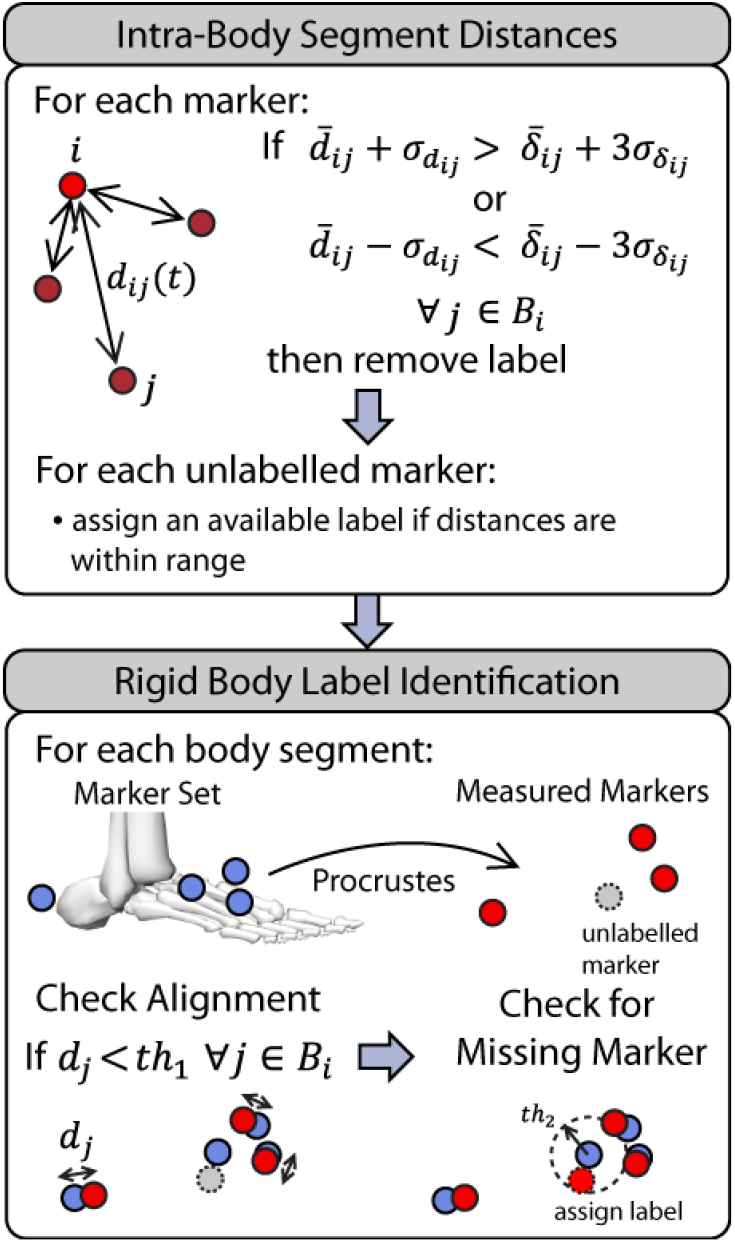
Overview of the marker set verification. First, the distances between a marker, *i*, and all other markers in its body segment, *B_i_*, were compared to the distances observed in the training data, *δ_ij_*. Then the marker coordinates defined in the marker set were used to attempt to identify unlabeled markers.

Finally, the marker set was used to attempt to assign labels for body segments where at least three markers were identified, but one or more were unlabelled. The local marker coordinates, defined in the OpenSim marker set, were scaled, rotated, and translated using a Procrustes analysis (Gower and Dijksterhuis, 2004) to align them with the measured markers. If a good alignment was achieved, indicated by the distances between the aligned marker set and measured markers all being less than a chosen threshold, *th*_1_, an attempt was made to locate the unlabelled marker. If there was an unlabelled marker within a threshold *th*_2_ of the expected position based on the aligned marker set coordinates, it was assigned the missing label. We used *th*_1_ = 3*cm* and *th*_2_ = 4*cm* based on the spacing of markers in our marker set.

### Analysis

We first explored the case where the algorithm is trained on pre-existing labelled motion capture data. The neural network was trained on the 100 participants in the Data Set A training set and the labelling algorithm was used to label the 42 participants in the test set. To assess performance, the predicted and ground truth labels for the markers were compared at each frame of data across all trials. The classification accuracy (micro-averaged F1 score), markers labelled incorrectly, and markers that should have been labelled but were not (falsely unlabelled) were calculated.

We then examined the performance for a different data set using simulated marker trajectories to train the neural network. The simulated marker trajectories for the Data Set B marker set were used to train the neural network. The algorithm was then used to label all trials from the nine participants in the Data Set B test set with the extraneous markers removed. Whereas Data Set A had previously been gap-filled, Data Set B provided a realistic test of the accuracy of the labelling procedure on raw marker data that contain occlusions, missing markers, marker data split into multiple trajectories, and movements outside of those in the training set. Metrics to quantify data quality were also calculated. We examined the sensitivity of the algorithm to the size of the training set by training the algorithm on subsets of simulated trajectories ranging from 10-100 participants.

In practice, it would be possible to add correctly labelled data into the training set of simulated data as the procedure is used over time. Therefore, we used transfer learning by starting with the algorithm trained on the simulated data from 100 participants and performed additional training on labelled data from Data Set B. We evaluated the effect of increasing the training set through transfer learning by performing additional training on a single trial from Data Set B and then for all trials from 1-9 participants in the training set of Data Set B. Each trained algorithm was used to label the nine participants in the test set. The addition of one (Sim+1) and nine (Sim+9) participants was investigated in more detail.

Lastly, the accuracy of the labelling procedure when extraneous markers are present was tested. The algorithm was trained on simulated marker trajectories used to label Data Set B with the extraneous markers retained. We calculated the same metrics as above as well as the percentage of markers that should not have been labelled but were (falsely labelled). Falsely labelled markers only exist if there are extraneous or ghost markers present in the data. The data of one participant with extraneous markers retained was used to update the neural network weights and the test set data was labelled. The same process was followed for updating based on nine participants with extraneous markers included. These results were compared with the automatic labelling with the extraneous markers retained performed in Vicon Nexus using the statically and dynamically calibrated skeleton models.

Five trials where the labelling accuracy for our algorithm and Nexus were within 1% of each other and were close to 90% were selected for manual correction in both Nexus and our GUI. The time to perform the correction for each trial was recorded.

## Results

### Data Set A – Training on Pre-Existing Data

Data set A had previously been gap filled and, as a result, had few occlusions and gaps in marker trajectories (Table 2). It included no extraneous physical markers and few trials where markers were missing for the entire trial.

**Table 2:** Description of the data sets used.

|  | Data Set A | Data Set B |
| --- | --- | --- |
| Number of Participants | 184 | 18 |
| Total Number of Trials | 2092 | 619 |
| Number of Frames per Trial | 512<br>(74-3645) | 1528<br>(137-12826) |
| Number of Labels | 45 | 62 |
| Number of Trajectories<sup>a</sup> | 45 (42-59) | 83 (59-533) |
| Number of Missing Markers<sup>b</sup> | 0 (0-3) | 0 (0-7) |
| Number of Extraneous Physical Markers | 0 (0-0) | 10 (7-23) |
| Marker Visibility (%)<sup>c</sup> | 100 (93.2-100) | 95.9 (79.0-99.8) |

<sup>a</sup>Number of trajectories indicates the number of ‘markers’ the marker trajectories were split into due to occlusions.
<sup>b</sup>Number of missing markers is a count of labels that were occluded or physically missing for the entire trial.
<sup>c</sup>Marker visibility is the percentage of frames where each marker is visible. Median values are presented with the range in parentheses

The accuracy of the marker labelling algorithm on the Data Set A test set was 99.6%. Across all data frames in the set, 0.1% of markers were assigned an incorrect label, and 0.3% of markers were not assigned a label (Table 3).

**Table 3:** Labelling performance on data sets a and b. For Data Set A, the algorithm was trained on motion capture data (MoCap). For Data Set B, the algorithm was trained on simulated trajectories (Sim) or simulated trajectories and updated with data from one (Sim+1) or nine participants (Sim+9). Data Set B was also automatically labelled using Vicon Nexus following static (Nexus-Stc) and dynamic (Nexus-Dyn) skeleton calibrations.

| Training Data | Accuracy (%) | Incorrect Label (%) | Falsely Unlabelled (%) | Falsely Labelled (%) |
| --- | --- | --- | --- | --- |
| Data Set A Test Set |  |  |  |  |
| MoCap | **99.6** | **0.1** | **0.3** | - |
| Data Set B Test Set (Extra Markers Removed) |  |  |  |  |
| Sim | 92.8 | 3.7 | 3.4 | - |
| Sim+1 | 97.1 | 1.4 | 1.5 | - |
| Sim+9 | **98.8** | **0.7** | **0.4** | - |
| Data Set B Test Set (Extra Markers Retained) |  |  |  |  |
| Sim | 83.5 | 5.8 | 8.0 | 2.8 |
| Sim+1 | 90.9 | 2.6 | 4.4 | 2.1 |
| Sim+9 | **95.6** | 1.0 | 1.8 | **1.6** |
| Nexus-Stc | 91.6 | 2.7 | 1.4 | 4.3 |
| Nexus-Dyn | **95.6** | **0.6** | **0.3** | 3.5 |

### Data Set B – Training on Simulated Trajectories

Data Set B comprised raw motion capture data before any labelling, filtering, or gap filling had occurred. Additionally, the movements performed caused marker occlusion at times. As a result, marker data were split into a greater number of trajectories and there were more occlusions than in Data Set A (Table 2).

The algorithm was trained on the simulated marker trajectories generated based on body kinematics of 100 participants from Data Set A. In the nine participants from the Data Set B test set, 92.8% of markers were labelled correctly across all frames when extraneous markers were removed (Table 3). 3.7% of markers were assigned an incorrect label and 3.4% were not labelled. Using transfer learning to update the neural network weights based on correctly labelled marker trajectories from one participant (Sim+1) improved the accuracy of the labelling procedure to 97.1%. Adding the labelled data of nine participants (Sim+9) further improved the accuracy to 98.8%.

In Data Set B, the participants performed the same athletic movements used in Data Set A as well as eight movements unknown to the neural network. When the neural network was trained on the simulated trajectories alone, 97.2% of markers for movements included in the training set were accurately labelled compared to 87.0% in previously unseen movements (Fig. 6A). However, updating the neural network weights by adding in labelled data from just one participant (Sim+1) increased the accuracy on these unknown movements to 95.0%, and further improvement to 98.2% was obtained by adding nine participants to the training data (Sim+9).

The number of participants used to train the algorithm had a greater effect on the accuracy on previously unknown movements than on the known movements contained in the training data (Fig. 5A). Adding labelled data to the simulated training set through transfer learning also had a larger effect on the accuracy for unknown compared to known movements (Fig. 5B). Adding a single trial increased the overall accuracy by 1% and adding all trials from one participant increased accuracy by 3.2%. Additional participants typically continued to improve accuracy by less than 1% per participant.

**Figure 5.**
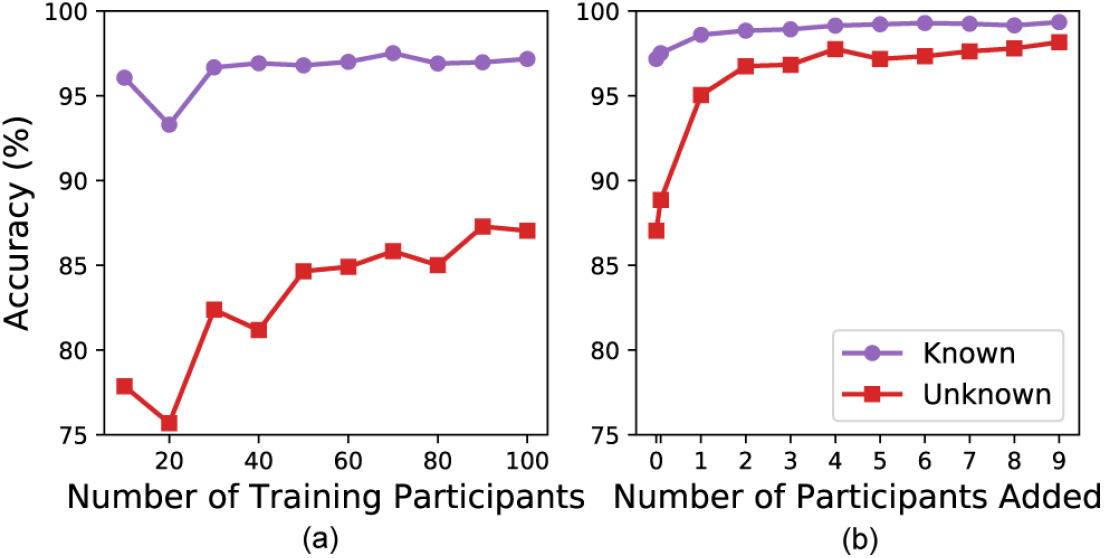
Accuracy of labelling the Data Set B test set with extraneous marker removed for known and unknown movements. (a) The effect of the number of participants included in the training set of simulated marker trajectories. (b) The effect of the number of participants used for transfer learning. A single trial from a single participant is also included.

When the extraneous markers from Data Set B were included in the input to the labelling algorithm, the overall accuracy was reduced to 83.5% (Table 3). When the algorithm was trained on only the simulated marker trajectories, the labelling accuracy was 87.4% for known movements and 78.2% for unknown movements (Fig. 6B). However, updating the neural network weights by performing additional training on labelled data including extraneous markers from one participant (Sim+1) increased the accuracy to 92.3% for known movements and 89.0% for unknown movements. Performing training on the data of nine participants (Sim+9) including extraneous markers improved the accuracy to 96.6% for known movements and 94.1% for unknown movements. Overall, the accuracy of the Sim+1 and Sim+9 algorithms had similar performance to the statically and dynamically calibrated models in Nexus, respectively. Nexus had a higher percentage of falsely labelled markers whereas the algorithm had more falsely unlabelled markers.

**Figure 6.**
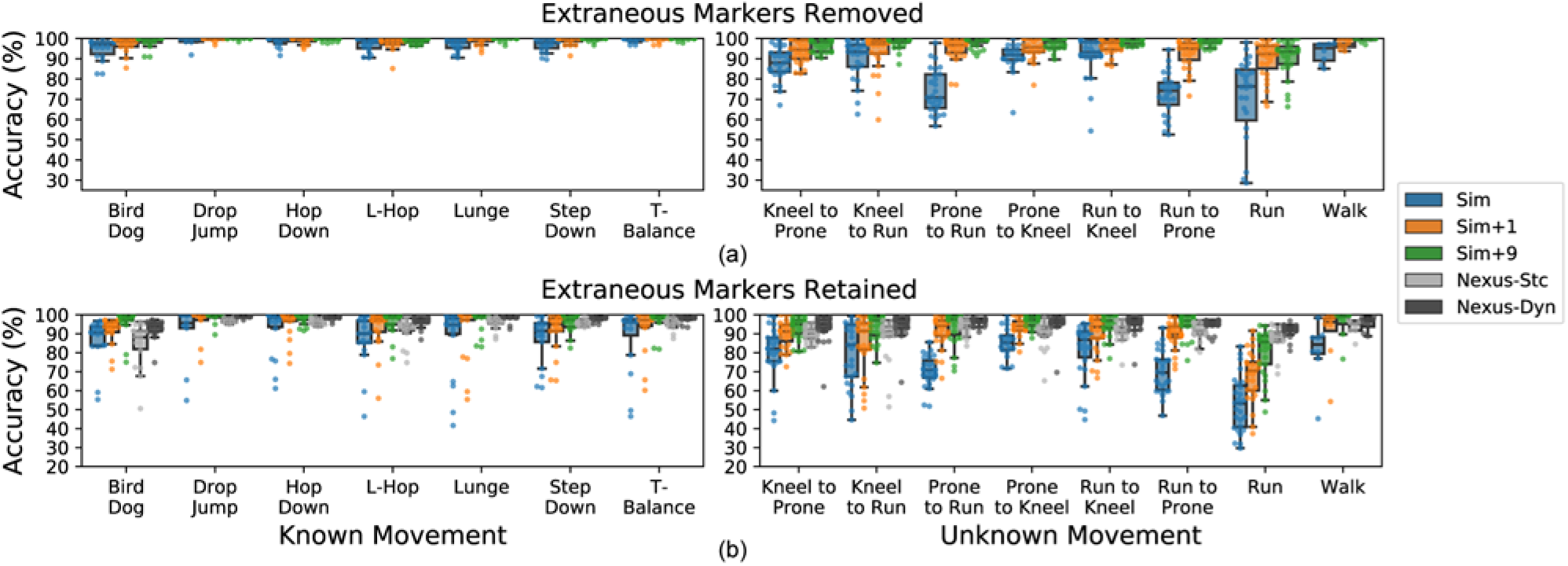
Accuracy of the labelling procedure on nine participants from Data Set B when trained on simulated trajectories (Sim) and updated using motion capture data from one (Sim+1) or nine participants (Sim+9) and accuracy of automatic labelling in Vicon Nexus following the static (Nexus-Stc) and dynamic (Nexus-Dyn) calibrations. Results are shown when extraneous markers are removed (a) or retained (b) and are separated by movement type.

The visibility of the markers, which indicates the amount of time the markers were not occluded, was associated with increased accuracy of the labelling algorithm (Fig. 7A). Whereas the number of markers missing from the entire trial had little effect when extraneous markers were removed, a greater number of missing markers negatively affected the accuracy when the extraneous markers were included (Fig. 7B). There was a similar relationship between these parameters and accuracy for the automatic labelling performed in Nexus using a static calibration (Fig. 7C).

**Figure 7.**
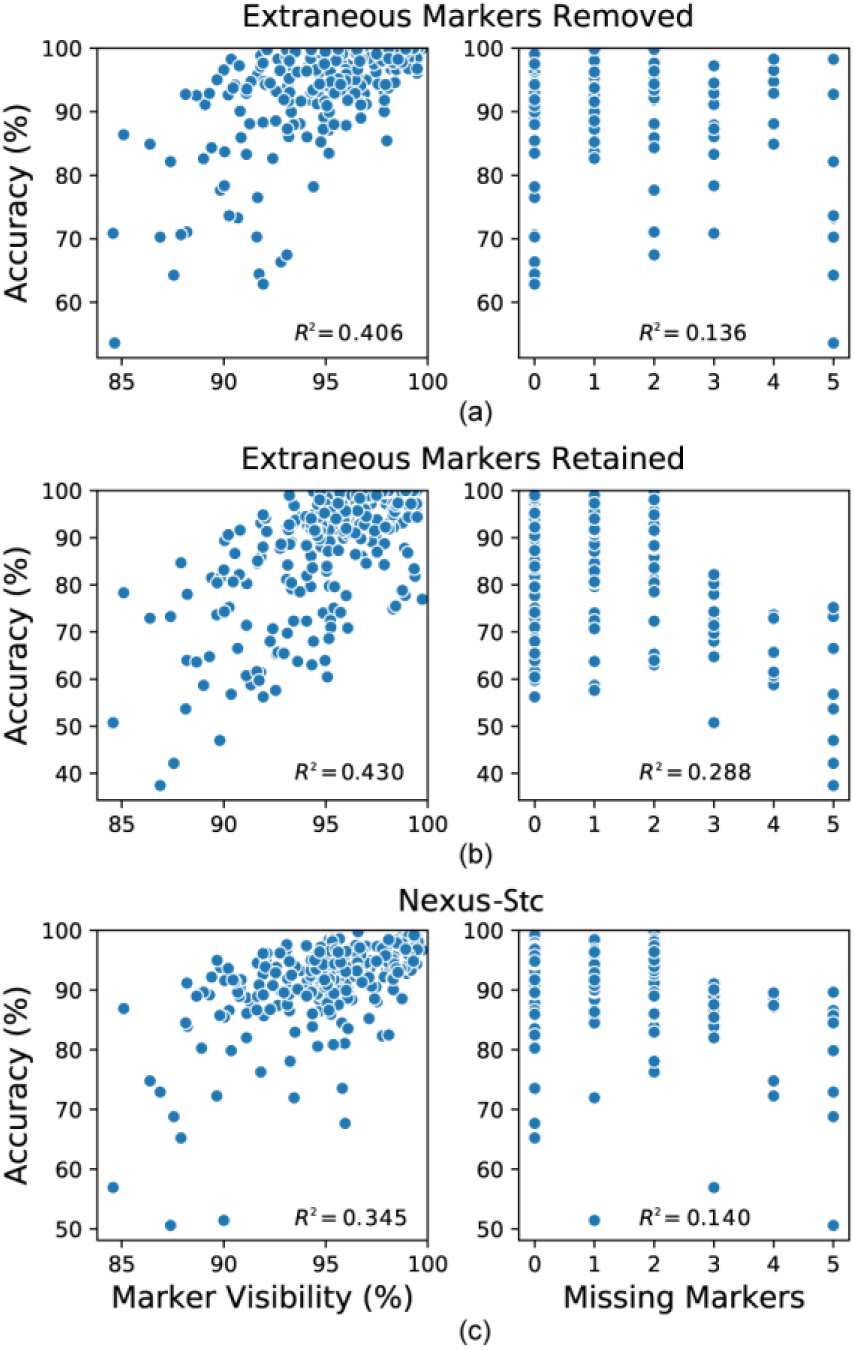
Associations of marker visibility and missing markers with labelling accuracy the Sim+1 models when extraneous markers were removed (a) or retained (b) and for the automatic labelling in Nexus following static calibrations (c). Missing markers indicated the number of labels missing from the entire trial. R^2^ values for linear regressions between the variables are shown.

The five selected trials took an average of 13.4 ± 6.7 minutes to manually correct in our GUI and 22.6 ± 23.9 minutes in Nexus. The difference in time was mainly due to one trial taking significantly longer (62 vs. 12 minutes) in Nexus; the times for the other trials were similar.

The median time to load and label a trial with our algorithm was 5.8 seconds (range 0.6-331.3s) with a mean processing speed of 254 ± 74 frames/s. The mean speed of the automatic labelling in Nexus was 130 ± 40 frames/s. In addition, a total of 32.8 minutes was spent to manually label static calibration trials for the nine participants in Nexus, which were used for the static calibration of the Nexus skeleton (Nexus-Stc). A total of 17.1 hours was spent to manually correct the labels in the functional range of motion trials in Nexus for the nine participants used for the dynamic calibrate of the Nexus skeleton (Nexus-Dyn).

### Graphical User Interface (GUI) and Source Code

To facilitate use of the algorithm, we created a graphical user interface that allows a user to import a .c3d file, run the automatic labelling routine, review the results, correct any errors, and export a labelled .c3d file (Fig. 8). The GUI provides error detection for unlabelled and duplicated markers. The marker visualization indicates the confidence in the predicted label based on the probability. The other options for visualization are to highlight unlabelled markers or to colour the markers based on their associated body segment. Mousing over a marker in the plot provides the marker number and current label. The GUI was tested on a Windows 10 desktop with an Intel^®^ Core™ i7-9700 CPU and 16GB of RAM. The data required to generate the simulated marker trajectories and source code are available at https://github.com/aclouthier/auto-marker-label.

**Figure 8.**
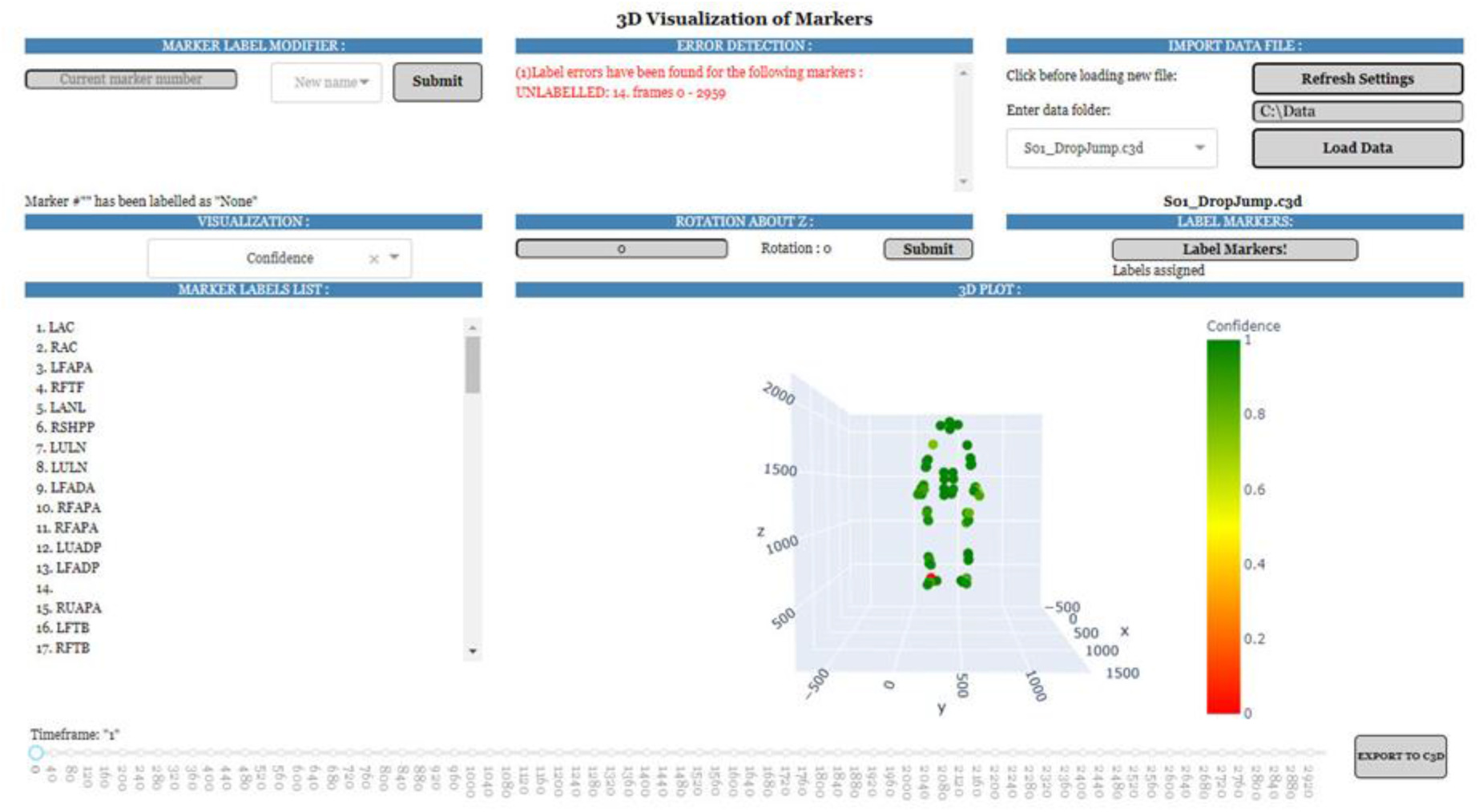
GUI that allows a user to run the automated marker labelling procedure on a .c3d file, make any necessary corrections, and export a labelled version. The visualization colours the markers based on the confidence in the predicted label.

## Discussion

We have developed an algorithm for automatically labelling motion capture markers using machine learning. The algorithm can be trained on measured or simulated marker trajectories. It is able to label movements outside of those contained in the training data with an opportunity to improve the accuracy by adding additional labelled motion capture data to the training set. There is no participant-specific initialization or calibration required. The algorithm is able to label markers in data that contain occlusions, ghost markers, missing markers, and extraneous markers. We have provided Python code for generating the simulated training data based on custom OpenSim marker sets, training the neural network, and running the labelling procedure as well as a GUI for performing the labelling and verifying the results.

The performance when training the algorithm on labelled motion capture data using Data Set A was excellent, with an accuracy of 99.6%. This Data Set represented close to optimal conditions as it contained high-quality data with few occlusions and no extraneous markers, no movements outside of those in the training set, and a large amount of labelled data to use for training. In contrast, we used Data Set B to examine a more realistic case where there were a larger number of occlusions, extraneous markers, movements outside of the training data, and no existing training set.

The marker labelling algorithm trained on simulated trajectories performed very well on the activities included in the simulated data, achieving an accuracy of 97.2% across all frames of data for nine participants in Data Set B. However, on previously unseen movements, the accuracy varied by task.

Walking trials and trials involving kneeling all had a mean accuracy greater than 80%, but the mean accuracies for running trials and trials transitioning between running and prone positions were between 70-80%. With the exception of the bird-dog, the training movements all involved upright postures, which may be the reason for this reduction in accuracy for prone positions. The accuracy was likely less for running movements because of the greater velocities and accelerations compared to the training movements as well as the participant entering and exiting the capture volume, reducing marker visibility at the start and end of the trial. We obtained continuous improvements in accuracy with each participant added to the training set through transfer learning. However, adding motion data from one participant in Data Set B improved the accuracy such that all movement types had mean accuracies greater than 90%. Therefore, in practice, this reduction in accuracy for movements outside of the simulated data can be largely eliminated once a few trials of the new movements have been labelled and added to the training set.

We have demonstrated that the algorithm can label markers in the presence of occlusions and extraneous markers. Data Set B included both temporarily occluded markers as well as up to five markers missing from the entire trial. A smaller marker visibility percentage was somewhat associated with lower accuracies (Fig. 7). The number of missing markers was associated with reduced accuracy only if the extraneous markers of the calibration wand, infrared camera, and additional anatomical markers were included. It was expected that decreased marker visibility and missing markers would negatively affect the performance and the correlations were similar to those resulting from the automatic labelling in the commercial software Nexus. When the algorithm was trained on simulated trajectories, the presence of extraneous markers in the trial decreased labelling accuracy, reducing it to 83.5% compared to 92.8% when extraneous markers were removed. A previously proposed approach similarly reported reduced accuracies when additional markers were present in the capture volume (Han et al., 2018). Updating the neural network weights based on the data from one participant, including extraneous markers, improved the accuracy to 90.9%, which was on par with the performance of the statically calibrated Nexus model at 91.6%. Our algorithm also offers the ability to further improve results by continuing to add data to the training set as more trials are labelled.

The processing speed of our algorithm was faster than that of the automatic labelling in Nexus. However, a significant advantage of our algorithm is not requiring a participant-specific calibration. This requires significant manual effort, especially for the dynamic calibration for which a functional range of motion trial must be labelled for each participant. If the algorithm is trained on existing motion capture data, then no manual intervention is required before labelling new data using the same marker set. If simulated marker trajectories are used in training, then some additional trials may need to be labelled, corrected, and added to the training set to attain the same accuracy as Nexus. However, these can be trials that would otherwise need to be corrected and once an acceptable accuracy is reached, no further manual effort is required for new participants. Additionally, we found that the time required to manually correct the labels using our GUI was similar to or less than when using Nexus, depending on the trial.

The presented labelling algorithm does have some limitations. The algorithm requires the participant to be facing in the positive x-direction and large deviations from this will reduce the accuracy. Ghorbani et al. (2019) proposed using principal component analysis to align the marker data during pre-processing; however, their data consisted of walking, jogging, sitting, and jumping tasks. This method will fail to align the marker data in postures that deviate from a standard posture, such as kneeling or having the arms raised above the head. Therefore, we have opted to perform this alignment manually, when it is necessary, and have included an option in our GUI to input the degrees to rotate about the vertical axis based on the marker visualization. The data are rotated back to their original orientation before being exported. Our simulated trajectory generation and marker set verification steps require a marker set to be defined in OpenSim. If our pipeline to generate simulated trajectories is used, the marker set must be based on the same OpenSim model used here. However, if existing data are used for training, any OpenSim model can be used, including the addition of external objects. Furthermore, the algorithm could be used without the marker set verification steps if the user does not wish to create the OpenSim marker set. This algorithm cannot be used for real-time marker labelling as currently implemented. However, it is intended for use in post-processing of data and is able to label more frames/s than Nexus and exceeds or is similar to previously reported speeds, which range from 4-251 frames/s (Maycock et al., 2015; Xia et al., 2017; Yu et al., 2007). The initial time to train the neural network on the simulated data is somewhat long due to the size of this data set (100 participants). It took 21 hours to train 10 epochs on an NVIDIA Titan RTX GPU. However, this only needs to be completed once for a given marker set and then no further initialization or calibration is required for individual participants. Furthermore, good accuracy can still be achieved using less training data, especially for movements contained in the training set (Fig. 5), which would reduce training time. Adding to the training set through transfer learning is also quicker, taking approximately one minute per trial. Finally, in the GUI, small gaps are interpolated using a cubic spline, but more advanced gap filling methods could be implemented in the future.

## Conclusion

The proposed algorithm is able to automatically label motion capture markers in the presence of occluded and extraneous markers. The algorithm can be trained on existing or simulated marker data. The accuracy of the algorithm is on par with commercial software and has the potential to improve as data are collected, labelled, and added to the training set. The code and data required to generate the simulated marker trajectories and train the algorithm and the GUI are open source, making the algorithm accessible for many motion capture users. This automated marker labelling algorithm has the potential to reduce the time and manual effort required to label motion capture data, especially for those who have limited access to commercial software.

